# Plant DNA polymerases alpha and delta mediate replication of geminiviruses

**DOI:** 10.1101/2020.07.20.212167

**Authors:** Mengshi Wu, Hua Wei, Huang Tan, Shaojun Pan, Qi Liu, Eduardo R Bejarano, Rosa Lozano-Durán

## Abstract

Geminiviruses are causal agents of devastating diseases in crops. Geminiviruses have circular single-stranded (ss)DNA genomes that are replicated in the nucleus of the infected plant cell through double-stranded (ds)DNA intermediates by the plant’s DNA replication machinery; which host DNA polymerase mediates geminiviral multiplication, however, has so far remained elusive. Here, we show that subunits of the nuclear replicative DNA polymerases α and δ physically interact with the geminivirus-encoded replication enhancer protein, C3, and are required for viral replication. Our results suggest that while DNA polymerase α is essential to generate the viral dsDNA intermediate, DNA polymerase δ mediates the synthesis of new copies of the geminiviral ssDNA genome, and that the virus-encoded C3 acts selectively recruiting DNA polymerase δ over ε to favour a productive replication.

## MAIN TEXT

Being obligate intracellular parasites, viruses rely on the host molecular machinery to replicate and spread. Geminiviruses are a family of plant viruses with circular single-stranded (ss) DNA genomes, causal agents of devastating diseases in crops worldwide (reviewed in Varma and Malathi, 2003). None of the geminivirus-encoded proteins is a DNA polymerase, and geminiviral replication, which occurs in the nuclei of infected cells, completely relies on the plant DNA replication machinery. In a first step, the viral ssDNA has to be converted into a double-stranded (ds) intermediate, which is then replicated by rolling-circle replication (RCR) and recombination-dependent replication (RDR), producing multiples copies of the original viral genome that are eventually encapsidated and can be transmitted by the insect vector (reviewed in Hanley-Bowdoin et al., 2013; Rizvi et al., 2015). Only one viral protein, the replication-associated protein (Rep), is required for the replication of viral DNA: Rep reprograms the cell cycle, recruits the host DNA replication machinery to the viral genome, and mediates nicking and rejoining events required for the initiation of replication and release of newly-synthesized molecules (reviewed in Hanley-Bowdoin et al., 2013; Rizvi et al., 2015). Another viral protein, C3, plays an ancillary role in viral DNA replication, acting as an enhancer in this process through an as-yet-unknown mechanism (Elmer et al., 1988; Hormuzdi and Bisaro, 1995; Morris et al., 1991; Stanley et al., 1992; Sung and Coutts, 1995; Sunter et al., 1990). A few host factors interacting with Rep or C3 and potentially required for geminiviral DNA replication, including the sliding clamp proliferating cell nuclear antigen (PCNA), the sliding clamp loader replication factor C (RFC), and the ssDNA binding protein replication protein A (RPA), have been described to date (Castillo et al., 2003; Luque et al., 2002; Singh et al., 2007; reviewed in Rizvi et al., 2015); a recent genetic screen has identified a number of factors required for geminivirus replication in yeast, which can act as a surrogate system (Li et al., 2020). Nevertheless, the plant DNA polymerase replicating the viral genome has so far remained elusive.

In order to identify host factors involved in the replication of geminiviral DNA, we performed a yeast two-hybrid screen using C3 from *Tomato yellow leaf curl virus* (TYLCV) as bait against a cDNA library from infected tomato plants (Rosas-Diaz et al., 2018). Interestingly, we found that C3 interacts with the N-terminal part of DNA polymerase α subunit 2 (SlPOLA2) (Figure 1A); this interaction was confirmed in yeast and in planta through co-immunoprecipitation (co-IP) and bimolecular fluorescence complementation (BiFC) assays, and could also be detected with POLA2 from the *Solanaceae* experimental host *Nicotiana benthamiana* (NbPOLA2) (Figure 1B-E; see Materials and Methods; Table S1). In addition, NbPOLA2 interacts with the C3 protein encoded by the geminiviruses *Beet curly top virus* (BCTV) and *Tomato golden mosaic virus* (TGMV) (Figure 1E), suggesting broad conservation of this interaction within the viral family. Of note, TYLCV induced the formation of nuclear speckles by POLA2 (Figure 1F and S1). Chromatin immunoprecipitation (ChIP) assays indicated that NbPOLA2 can bind the viral genome in a C3-independent manner (Figure 1G and S2). Knocking-down *NbPOLA2* by *Tobacco rattle virus* (TRV)-mediated virus-induced gene silencing (VIGS) in *N. benthamiana* rendered plants with reduced height and thicker leaves (Figure S3), and, strikingly, although it did not affect Agrobacterium-mediated transient transformation (Figure S4), it almost completely impaired local TYLCV replication and systemic infection (Figure 1H, I), indicating an essential role of POLA2/DNA polymerase α in the replication of the viral genome. A similar effect of *NbPOLA2* silencing was observed on BCTV replication (Figure S5A), suggesting that the role of this polymerase is likely conserved across different geminivirus species.

**Figure 1.**
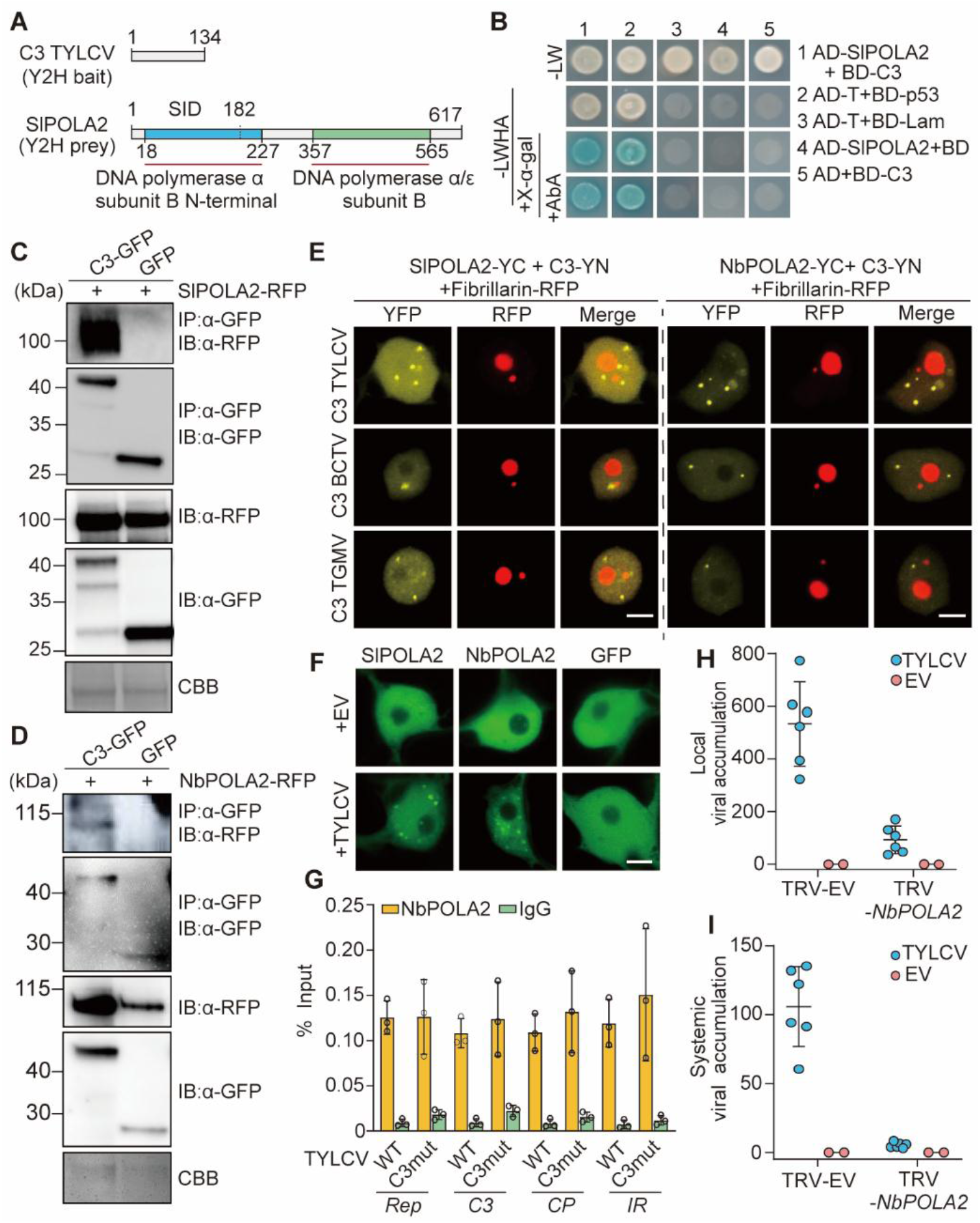
The DNA polymerase α subunit POLA2 interacts with the geminivirus-encoded C3 protein and is required for geminiviral replication. **A.** Schematic representation of bait (C3) and prey (SlPOLA2) isolated from the Y2H screen. Amino acid residues are indicated. SID: Selected interaction domain. **B.** C3 and POLA2 interact in yeast. AD: Activation domain. BD: Binding domain. **C, D.** C3-GFP co-immunoprecipitates SlPOLA2-RFP (C) and NbPOLA2 (D) upon transient expression in *N. benthamiana*. CBB: Commassie brilliant blue. **E.** SlPOLA2 and NbPOLA2 interact with C3 from TYLCV, BCTV, and TGMV in BiFC assays upon transient expression in *N. benthamiana*. Fibrillarin-RFP marks the nucleolus and the Cajal body. Scale bar: 5 µm. **F.** Nuclear distribution of transiently expressed SlPOLA2-GFP, NbPOLA2-GFP, and free GFP in the absence (empty vector, EV) or presence of TYLCV in *N. benthamiana*. Scale bar: 5 µm. Additional images are shown in Figure S1. **G.** NbPOLA2 binds the TYLCV genome in ChIP assays. The location of primers used for different genomic regions is shown in Figure S2. Error bars indicate SD of n=3 independent biological replicates. **H, I.** Viral accumulation in local (H; 3 days post-inoculation) or systemic (I; 14 days post-inoculation) TYLCV infections in *POLA2*-silenced (TRV-NbPOLA2) or control (TRV) *N. benthamiana* plants measured by qPCR. Plants inoculated with the empty vector (EV) are used as negative control. Error bars represent SD with n=6 independent biological replicates. The 25S ribosomal DNA interspacer (*ITS*) was used as reference gene; values are presented relative to *ITS*. All experiments were repeated at least three times with similar results.

In eukaryotes, DNA polymerase α primes DNA replication, while DNA polymerases δ and ε act as the main processive replicative polymerases. These three replicative polymerases are assembled into a large complex termed replisome, which contains all proteins required for DNA replication, including PCNA, RFC, and RPA (reviewed in Pedroza-Garcia et al., 2019). In yeast, DNA polymerase δ elongates the RNA/DNA primers produced by DNA polymerase α on both strands (Garbacz et al., 2018) and, according to the generally accepted model, then synthesizes the lagging strand, with DNA polymerase ε synthesizing the leading strand (Pursell et al., 2007); additionally, DNA polymerase δ is believed to perform initiation and termination of replication in both strands (Zhou et al., 2019). Recently, an alternative model has been proposed, according to which DNA polymerase δ would replicate both strands of the DNA, and the switch to DNA polymerase ε would only occur following replication errors (Johnson et al., 2015). The DNA polymerase δ subunit NbPOLD2, but not the DNA polymerase ε subunit NbPOLE2/DPB2, associates with C3 from TYLCV in co-IP experiments, and interacts with C3 from TYLCV, BCTV, and TGMV in BiFC assays (Figure 2A-D; see Materials and Methods; Table S1). Intriguingly, co-expression of NbPOLD2 and geminiviral C3 led to a dramatic change in the morphology of the fibrillarin-positive nucleolus/Cajal body (Figure 2A). Both NbPOLD2 and NbPOLE2/DPB2, however, can bind the viral genome in ChIP assays (Figure 2E, F and Figure S6). Lack of C3 in a TYLCV C3 null mutant enhances binding of POLE2 to the viral DNA, while increasing variation in the binding of POLD2 (Figure 2E, F). With the aim to decipher whether DNA polymerase α acts in concert with DNA polymerases δ and/or ε in the replication of the viral DNA, we silenced the corresponding subunits *NbPOLD2* and *NbPOLE2/DPB2* by VIGS in *N. benthamiana*, and tested the capacity of TYLCV to replicate in local infection assays and to infect systemically. Silencing of either subunit results in a distinct developmental phenotype, with plants of smaller size with leaves of abnormal shape (Figure S3) but, while knocking-down *POLD2* impaired TYLCV DNA replication in local infection assays (Figure 2G), knocking-down *POLE2* enhanced viral accumulation both locally and systemically (Figure 2H, I). The same effect could be observed for BCTV, indicating that the role of DNA polymerases δ and ε is likely conserved across the geminivirus family (Figure S5B, C). Silencing of *NbPOLA2* led to a mild increase in the expression of *NbPOLD2* and *NbPOLE2/DPB2*; silencing of *NbPOLD2* or *NbPOLE2* did not trigger an up-regulation of the reciprocal subunit, ruling out compensatory effects (Figure S7). Taken together, these results indicate that DNA pol δ, but not DNA pol ε, is required, together with DNA pol α, for replication of the geminiviral genome.

**Figure 2.**
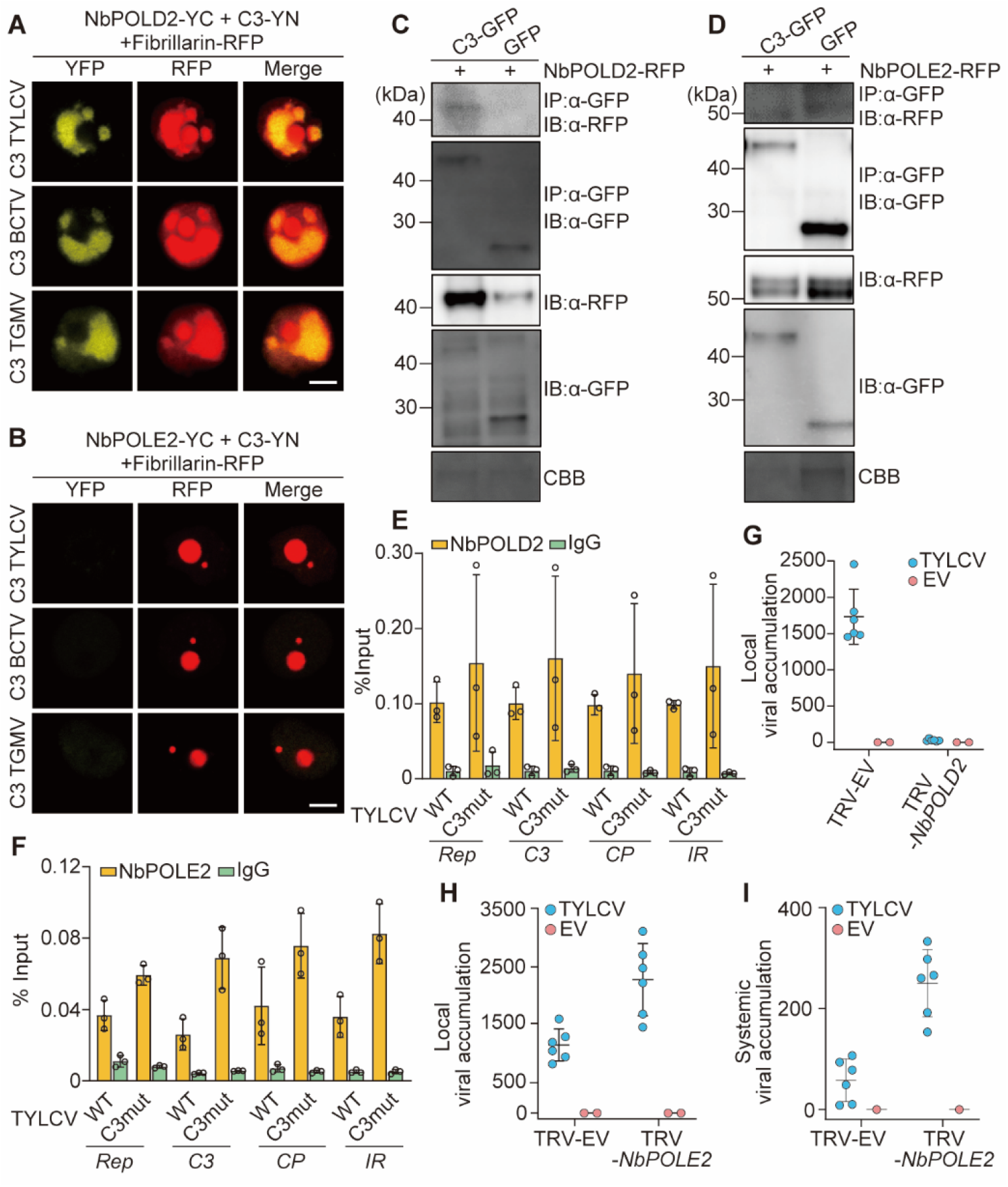
The DNA polymerase δ subunit POLD2, but not the DNA polymerase ε subunit POLE2, interacts with the geminivirus-encoded C3 protein and is required for geminiviral replication. **A, B.** NbPOLD2 (A), but not NbPOLE2 (B), interacts with C3 from TYLCV, BCTV, and TGMV in BiFC assays upon transient expression in *N. benthamiana*. Fibrillarin-RFP marks the nucleolus and the Cajal body. Scale bar: 5 µm. **C, D.** C3-GFP co-immunoprecipitates NbPOLD2-RFP (C), but not NbPOLE2-RFP (D), upon transient expression in *N. benthamiana*. CBB: Commassie brilliant blue. **E, F.** NbPOLD2 (E) and NbPOLE2 (F) bind the TYLCV genome in ChIP assays. The location of primers used for different genomic regions is shown in Figure S2. Error bars indicate SD of n=3 independent biological replicates. **G, H.** Viral accumulation in local TYLCV infections (3 days post-inoculation) in *POLD2-*silenced (TRV-NbPOLD2) (G), *POLE2*-silenced (TRV-NbPOLE2) (H), or control (TRV) *N. benthamiana* plants measured by qPCR. Plants inoculated with the empty vector (EV) are used as negative control. Error bars represent SD with n=6 independent biological replicates. **I.** Viral accumulation in systemic TYLCV infections (14 days post-inoculation) in *POLE2*-silenced (TRV-NbPOLE2) or control (TRV) *N. benthamiana* plants measured by qPCR. Plants inoculated with the empty vector (EV) are used as negative control. Error bars represent SD with n=6 independent biological replicates. The 25S ribosomal DNA interspacer (*ITS*) was used as reference gene; values are presented relative to *ITS*.

We next used two-step anchored qPCR (Rodriguez-Negrete et al., 2014) to quantify the relative accumulation of viral ssDNA (viral strand) and dsDNA intermediate (as complementary strand) (Figure 3A) in local infection assays with TYLCV WT or a C3 null mutant in *N. benthamiana* plants in which *POLA2*, *POLD2*, or *POLE2* have been silenced by VIGS (Figure 3B, C). In agreement with previous results, the lack of C3 resulted in a decrease in the accumulation of both strands of the viral DNA, consistent with the role of this viral protein as a replication enhancer (Figure 3B, C). Strikingly, silencing of *POLA2* impaired the accumulation of the viral complementary strand (Figure 3B), hence compromising the subsequent production of the viral strand (Figure 3C); silencing of *POLD2* did not affect the synthesis of the viral complementary strand, but interfered with the downstream accumulation of viral ssDNA (Figure 3B, C). As previously observed (Figure 2H), the lower levels of *POLE2* led to an increased accumulation of viral DNA, an effect that could be observed on both viral strands (Figure 3B, C). Taken together, these results suggest that DNA polymerase α is essential for the initial synthesis of the viral complementary strand, and therefore for the generation of the dsDNA replicative intermediate, while DNA polymerase δ, alone or in combination with α, is required for the following RCR. Interestingly, TYLCV has been recently proven to replicate in the insect vector in a DNA polymerase δ-dependent manner (He et al., 2020), which raises the idea that the mechanisms replicating geminiviruses might be conserved between the animal and plant kingdoms.

**Figure 3.**
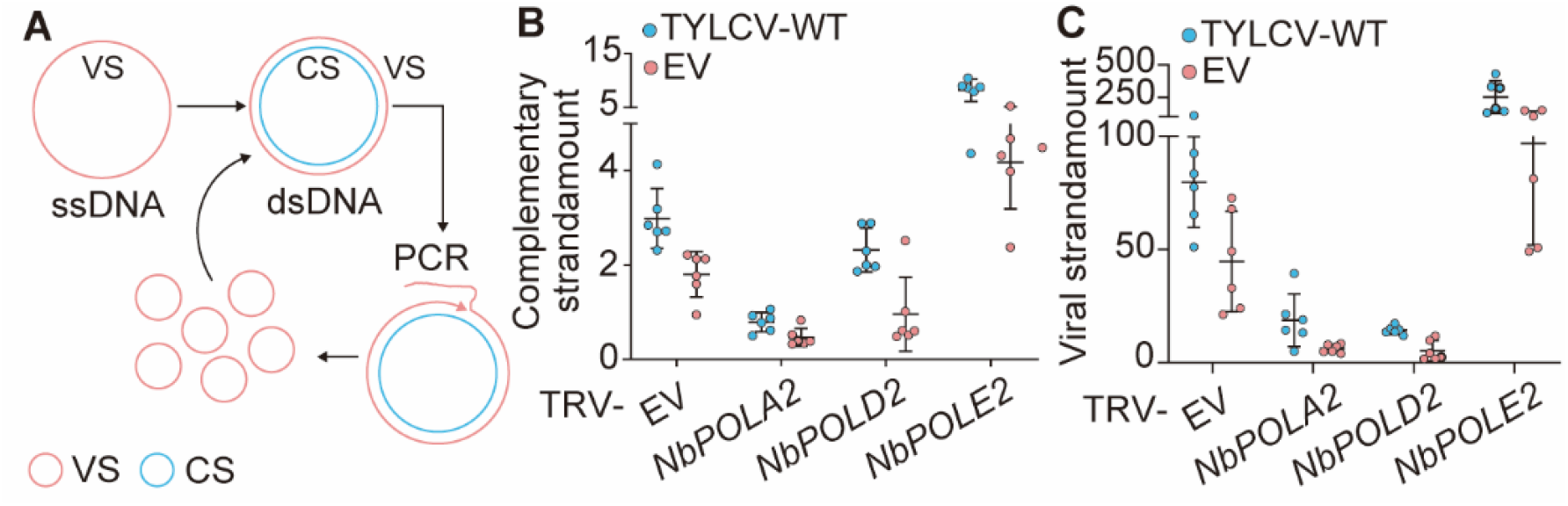
Effect of silencing *POLA2*, *POLD2*, and *POLE2* on the accumulation of viral and complementary DNA strands. **A.** Schematic representation of the viral DNA forms during the infection. VS: viral strand; CS: complementary strand; ssDNA: single-stranded DNA; dsDNA: double-stranded DNA. **B, C.** Accumulation of complementary strand (B) and viral strand (C) during local TYLCV infections in *POLA2*-silenced (TRV-NbPOLA2), *POLD2*-silenced (TRV-NbPOLD2), *POLE2*-silenced (TRV-NbPOLE2), or empty vector control (EV) *N. benthamiana* plants, measured by qPCR at 3 days post-inoculation. Error bars represent SD with n=6 independent biological replicates. The 25S ribosomal DNA interspacer (*ITS*) was used as reference gene; values are presented relative to *ITS*.

The finding that both DNA polymerases δ and ε can associate to the geminiviral genome, but δ is required for viral replication while ε seems to exert a negative effect that is released by its silencing, suggests that both DNA polymerases may compete for binding to the viral DNA with opposite outcomes, since only DNA polymerase δ leads to replication. The observations that i) C3 can physically interact with NbPOLD2, and ii) the non-productive binding of NbPOLE2 to the viral DNA is increased in the absence of C3, hint at the possibility that C3 might mediate the selective recruitment of DNA polymerase δ over ε. Supporting this hypothesis, silencing of *POLE2* increases DNA replication of a C3 null mutant TYLCV to wild type-like levels in control plants (Figure 3C). In order to further test this idea, we inoculated *N. benthamiana POLE2*-silenced or control plants with a TYLCV C3 null mutant, and evaluated the capacity of this mutant virus to establish a systemic infection. In systemic tissues of control plants, the mutant virus accumulated to very low levels, and caused only mild symptoms (Figure 4); nevertheless, upon *POLE2* silencing, both viral accumulation and symptom development were dramatically increased (Figure 4A-C). The ability of *POLE2* silencing to complement the lack of C3 strengthens the idea that one of the functions of this viral protein is to counter the negative effect of DNA polymerase ε on viral replication.

**Figure 4.**
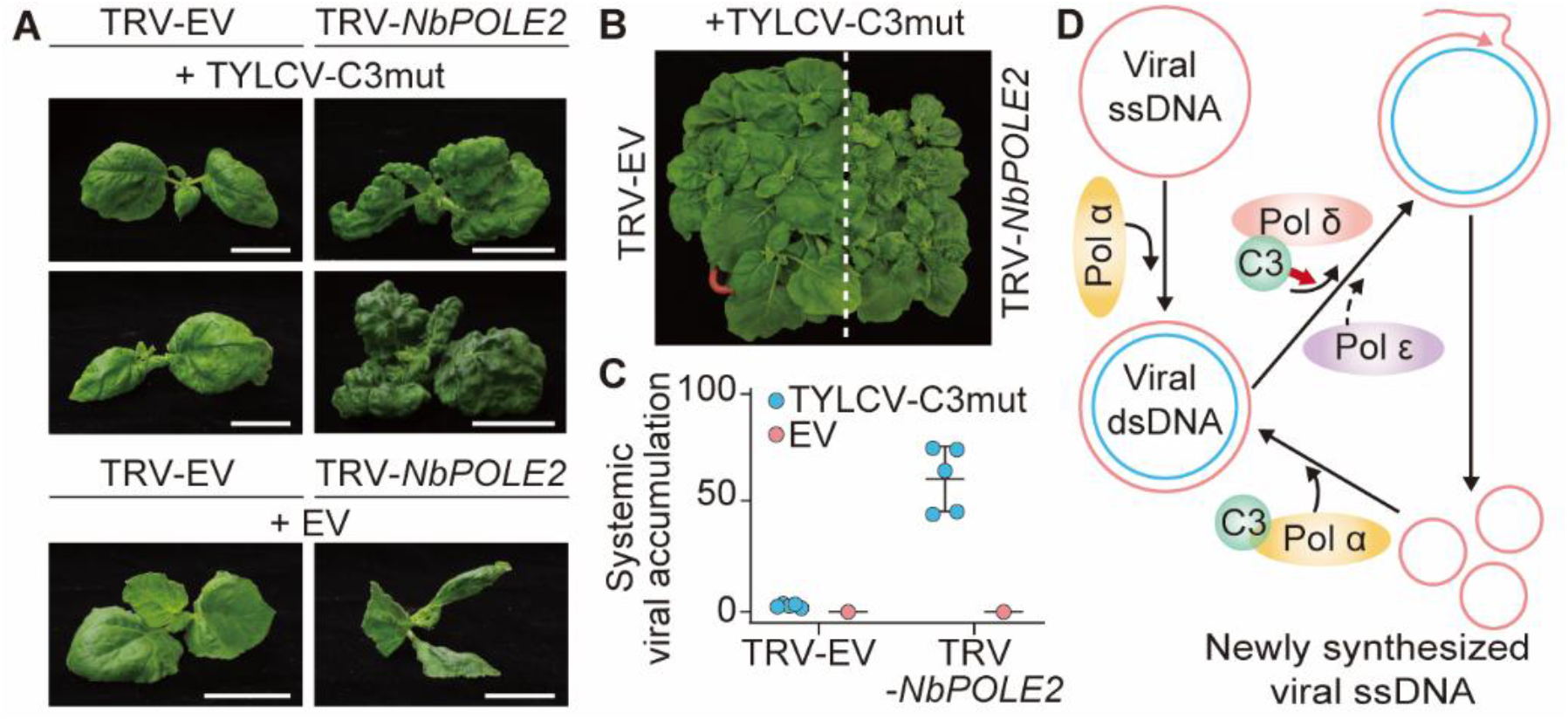
Silencing *POLE2* enables systemic infection of a C3 null mutant geminivirus. **A, B**. Symptoms of *POLE2*-silenced (TRV-NbPOLE2; right) or control (TRV-EV; left) *N. benthamiana* plants inoculated with a C3 null mutant virus at 28 days post-inoculation. EV: empty vector control. **C**. Viral accumulation in systemic TYLCV infections (28 days post-inoculation) in *POLE2-*silenced (TRV-NbPOLE2) or control (TRV-EV) *N. benthamiana* plants measured by qPCR. Plants inoculated with the empty vector (EV) are used as negative control. Error bars represent SD with n=6 independent biological replicates. The 25S ribosomal DNA interspacer (*ITS*) was used as reference gene; values are presented relative to *ITS*. **D.** Hypothetical model of the role of DNA polymerases α and δ in the replication of the geminiviral genome. DNA polymerase α is required to convert the viral ssDNA genome to the dsDNA replicative intermediate, which is then replicated by DNA polymerase δ to produce new viral ssDNA. The virus-encoded C3 protein interacts with DNA polymerase α (POLA2) and DNA polymerase δ (POLD2), and selectively recruits the latter over the non-productive DNA polymerase ε.

In summary, our results identify DNA polymerases α and δ as the DNA polymerases mediating replication of geminiviruses in their host plants, and suggest a model in which the geminivirus-encoded replication enhancer C3 acts selectively recruiting DNA polymerase δ over the non-productive DNA polymerase ε (Figure 4D). In yeast, DNA polymerase δ acts replicating the leading strand during double-strand break repair, a process in which it is error-prone (Deem et al., 2011; Hicks et al., 2010); during the replication of the geminiviral genome, a similar decrease in fidelity might explain the high mutation rate of these viruses. Geminiviruses belong to the CRESS DNA viruses, a virus phylum that encompasses Rep-encoding ssDNA viruses with a likely common ancestor infecting organisms from different kingdoms of life, including animals, plants, and fungi (Krupovic et al., 2020; Zhao et al., 2019); since CRESS DNA viruses are expected to display similar strategies for the replication of their genomes, this work could have a broad impact on host-virus interactions.

## ACKNOWLEDGEMENTS

The authors thank all members of Rosa Lozano-Duran’s lab and Alberto Macho’s lab for fruitful discussions, Xinyu Jian, Aurora Luque, and the PSC Cell Biology Facility for technical assistance, and Alberto P Macho, Chaonan Shi, and Laura Medina-Puche for critical reading of the manuscript. This work was supported by the Strategic Priority Research Program of the Chinese Academy of Sciences (Grant No. XDB27040206) and the Shanghai Center for Plant Stress Biology from the Chinese Academy of Sciences.

## SUPPLEMENTARY MATERIALS

Materials and Methods

Figures S1-S7

Tables S1-S3

## Notes

### Competing Interest Statement

The authors have declared no competing interest.

