## Supplementary materials for "Plant DNA polymerases alpha and delta mediate replication of geminiviruses"

**This file includes:**

Materials and Methods
Figs. S1 to S7
Tables S1 to S3

### MATERIALS AND METHODS

#### Plant materials

*N. benthamiana* plants were grown in a controlled growth chamber under long day conditions (LD, 16 h light/8 h dark) at 25°C.

#### Plasmid construction

Plasmids and primers used for cloning are summarized in Table S2 and Table S3. The TYLCV clone used as template is AJ489258 (GeneBank). pGTQL1211YN and pGTQL1221YC are described in Lu et al., 2010; vectors from the pGWB series are described in Nakagawa et al., 2007a, 2007b. DNA fragments clones into the pENTR$^{TM}$/D-TOPO entry vector or the pDNOR-zeo entry vector (Thermo Scientific) were recombined into the corresponding destination vectors through the Gateway LR reaction (Thermo Scientific).

#### Y2H assay

Yeast two-hybrid assays were performed following the Matchmaker Yeast Two-Hybrid User Manual (Clontech).

#### Identification and selection of *Nicotiana benthamiana* orthologue genes encoding DNA polymerase subunits

Orthologues of Arabidopsis POLA2, POLD2, and POLE2 proteins in *N. benthamiana* were identified by BLAST (Table S1). The proteins and coding genes used in this work were selected among the corresponding orthologues based on their gene expression in TYLCV-infected *N. bethamiana* samples (Wu et al., 2019; Table S1).

#### Local and systemic viral infections

Local and systemic viral infection assays were done as described in Medina-Puche et al., 2019. In brief, *Agrobacterium* cells carrying the wild-type TYLCV (TYLCV-WT), a C3 null mutant (TYLCV-C3mut), or the wild-type BCTV infectious clones (Table S2), or an empty vector (EV) as control, were liquid-cultured in LB with appropriate antibiotics overnight. Bacterial cultures were centrifuged at 4,000 g for 10 min and re-suspended in infiltration buffer (10 mM $MgCl_2$, 10 mM MES pH 5.6, 150 µM acetosyringone). After a 4-hour incubation in the dark, bacterial cultures were used to infiltrate the underside of leaves of 4-week-old *N. benthamiana* plants (for local infection assays) or inject on the stems of 3-week-old *N. benthamiana* plants (for systemic infection assays).

**Virus-induced gene silencing (VIGS)**

Tobacco rattle virus (TRV)-mediated virus-induced gene silencing (VIGS) assays were performed as described in Medina-Puche et al., 2019. Briefly, *Agrobacterium* cells carrying pTRV1 and pTRV2-based constructs (Table S2) were grown in LB medium overnight with appropriate antibiotics. Cultures were re-suspended in the infiltration buffer (10 mM $MgCl_2$, 10 mM MES pH 5.6, 150 µM acetosyringone) and incubated at room temperature for 4 h in the dark. Mixed cell cultures were used to inoculate two-week-old *N. benthamiana* plants. Two weeks later, plants were used for local infection assays. For systemic infection assays, *Agrobacterium* cells carrying the virus infectious clones were co-infiltrated with pTRV1 and pTRV2-based constructs.

**Quantitative real-time PCR (qPCR) and Reverse Transcription PCR (RT-qPCR)**

To determine viral accumulation, total DNA was extracted from *N. benthamiana* leaves (from infiltrated leaves in local infection assays and from apical leaves in systemic infection assays) using the CTAB method (Minas et al., 2011). Quantitative real-time PCR (qPCR) was performed with primers to amplify *Rep* (Table S3). The 25S ribosomal DNA interspacer (*ITS*) was used as reference gene (Table S3).

The quantification of viral strand and complementary strand in local infections was performed following (Rodriguez-Negrete et al., 2014).

To detect gene expression in *N. benthamiana*, total RNA was extracted from leaves by using Plant RNA kit (OMEGA Bio-tek). cDNA was synthesized using the iScript™ gDNA clear cDNA Synthesis Kit (Bio-Rad) according to the manufacturer's instructions. *NbActin* was used as reference gene. qPCR and RT-qPCR were performed in a BioRad CFX96 real-time system with Hieff™ qPCR SYBR Green Master Mix (Yeason). The reactions were done as follows: 3 min at 95 °C, 40 cycles consisting of 15 s at 95 °C, 30 s at 60 °C. Primers used are described in Table S3.

**Chromatin immunoprecipitation (ChIP) assay**

ChIP assay was performed as described previously (Nie et al., 2019). *Agrobacterium* clones carrying the binary vectors to express *NbPOLA2*-, *NbPOLD2*- or *NbPOLE2-GFP* (Table S2) were co-infiltrated with those carrying the TYLCV or TYLCV-C3mut infectious clones in *N. benthamiana* leaves. The infiltrated tissue was collected and crosslinked with 1% formaldehyde in 1xPBS buffer at 2 dpi. Then the tissue was ground in liquid nitrogen and re-suspended in Honda buffer (2.5% Ficoll 400, 5% Dextran T40, 0.4 M Sucrose, 25 mM Tris pH 7.4, 10 mM $MgCl_2$, 0.035% βmercaptoethanol, 1% Protease Inhibitor Cocktail (Sigma)), homogenized, and filtered through Miracloth (Millipore). 0.5% Triton X-100 was added to the supernatant, and the mixture was kept on ice for 15 min. After spinning at 2,000×g for 20 min at 4°C, the pellet was re-suspended in Honda buffer with 0.1% Triton X-100 and spun at 2,000×g for 10 min at 4°C. Isolated nuclei were re-suspended in 500 µL of Nuclei Lysis buffer and sonicated by BioruptorTM UCD-200 sonicator (diagenode) for 45 min. Following centrifugation at max speed for 5 min at 4°C, the supernatant was collected and used for input and immunoprecipitation. After adding 9 volumes of ChIP dilution buffer to the supernatant, this was pre-cleared with 10 µl of Dynabeads Protein G (Invitrogen) for 1 h at 4°C. After removing the beads from the mixture, the supernatant was incubated with anti-GFP antibody (Abcam, ab290), or anti-IgG antibody (Sigma, I5006) overnight at 4°C. The following day, after adding 20 µl of Dynabeads Protein G, the mixture was incubated for 2 h at 4°C. Beads were sequentially washed with 1 ml of the following buffers: twice Low Salt Wash buffer (150 mM NaCl, 0.1% SDS, 1% Triton x-100, 2 mM EDTA, 20 mM Tris pH 8.0), once High Salt Wash buffer (500 mM NaCl, 0.1% SDS, 1% Triton x-100, 2 mM EDTA, 20 mM Tris pH 8.0), once LiCl wash buffer (250 mM LiCl, 1% Igepal, 1% Sodium Deoxycholate, 1 mM EDTA, 10 mM Tris pH 8.0), twice TE buffer (10 mM Tris pH 8.0, 1 mM EDTA). Immunocomplexes were eluted with 150 µl of Elution buffer (1% SDS, 0.1 M $NaHCO_3$) at 65°C for 30 min. After reverse crosslinking, 10 µl of 0.5 M EDTA, 20 µl of 1 M Tris pH 6.5 and 1 µl of proteinase K (Invitrogen, 20 mg/mL) were added to each sample, which was incubated at 45 °C for 2 h. DNA was then purified using QIAquick PCR Purification Kit (QIAGEN, Cat. No. 28106). ChIP products were diluted into 200 µL of dd$H_2O$, and inputs were diluted at the ratio of 1:100, and analyzed by qPCR. The primers used in this experiment are listed in Table S3.

**Protein extraction and Co-immunoprecipitation (CoIP) assays**

*Agrobacterium* cells carrying the appropriate constructs were infiltrated *in N. benthamiana* leaves and collected at 2 dpi. After grinding the agroinfiltrated tissue in liquid nitrogen, nuclei were extracted as in ChIP assay, and then subjected to protein extraction and co-immunoprecipitation assays with GFP-Trap beads (Chromotek, Germany; Smart Lifesciences, SA070005) as described in (Rosas-Diaz et al., 2018). The antibodies used are as follows: anti-GFP (Abiocode, M0802-3a), anti-Red (Chromotek, 5F8), anti-Mouse IgG (Sigma, A2554), and anti-Rat IgG (Abcam, ab7097).

### Protein subcellular localization

For subcellular localization, GFP- or RFP-tagged proteins were transiently expressed in *N. benthamiana* leaves and imaged with a Leica TCS SMD confocal microscope using the preset settings for GFP (Ex: 488 nm, Em: 500-550 nm) and RFP (Ex: 554 nm, Em: 570-620 nm).

### Bimolecular Fluorescence Complementation (BiFC)

Bimolecular fluorescence complementation (BiFC) assays were performed in *N. benthamiana* leaves as described in Lu et al., 2010. *Agrobacterium* cells carrying the appropriate BiFC clones and RFP-Fibrillarin were infiltrated on 4-week-old *N. benthamiana* plants with 1 mL needle-less syringe. Imaging was performed two days later under a Leica TCS SMD confocal microscope by using the preset sequential scan settings for YFP (Ex: 514 nm, Em: 525-575 nm) and for RFP (Ex: 554 nm, Em: 570-620 nm).

**SUPPLEMENTARY FIGURES**

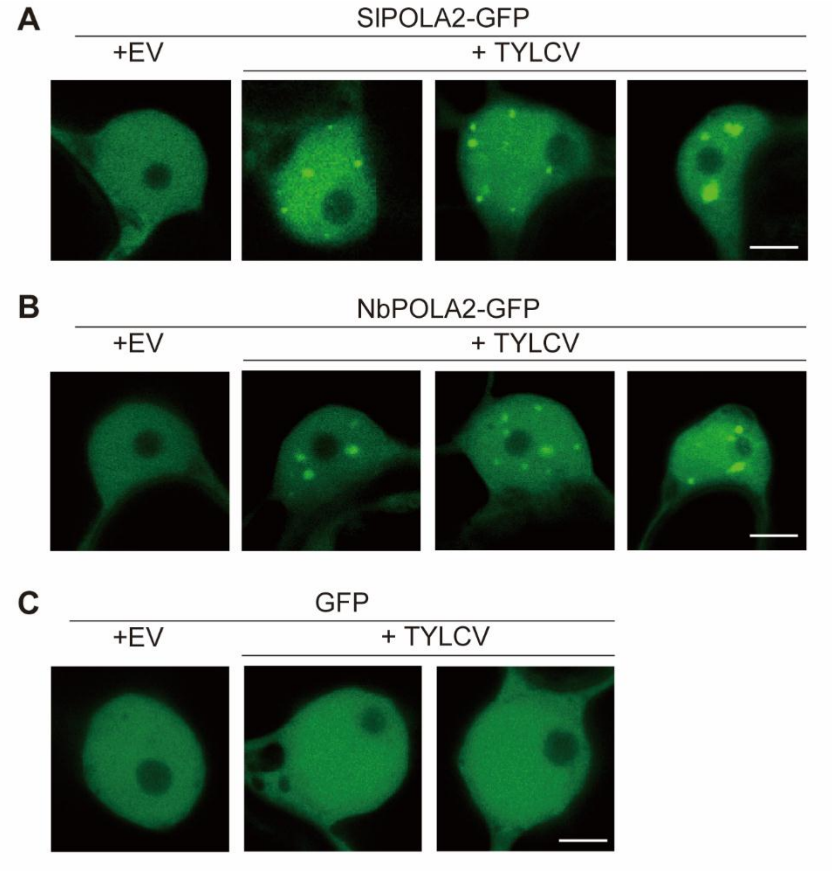


**Figure S1. Nuclear distribution of transiently expressed SlPOLA2-GFP (A), NbPOLA2-GFP (B), and free GFP (C) in the absence (empty vector, EV) or presence of TYLCV in *N. benthamiana*.** Scale bar: 5 µm. Additional images can be found in Fig. 1F. This experiment was repeated twice with similar results.

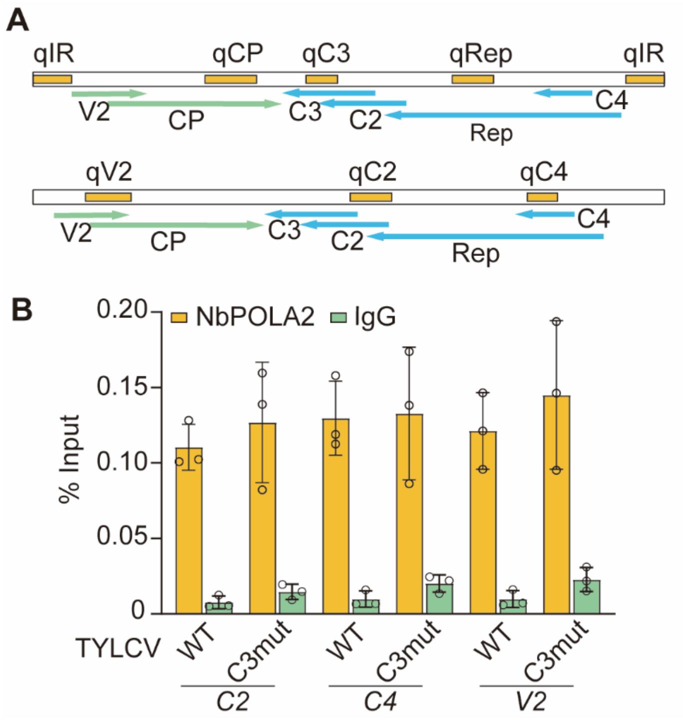


**Figure S2. Genomic location of the viral regions amplified in the ChIP assays (in yellow) (A), and binding of NbPOLA2 to the C2, C4, and V2 regions (B).** In A, viral genes are depicted as arrows; genes in the viral strand are coloured in light green; genes in the complementary strand are coloured in blue. In B, error bars represent SD with n=3 independent biological replicates. This experiment was repeated twice with similar results.

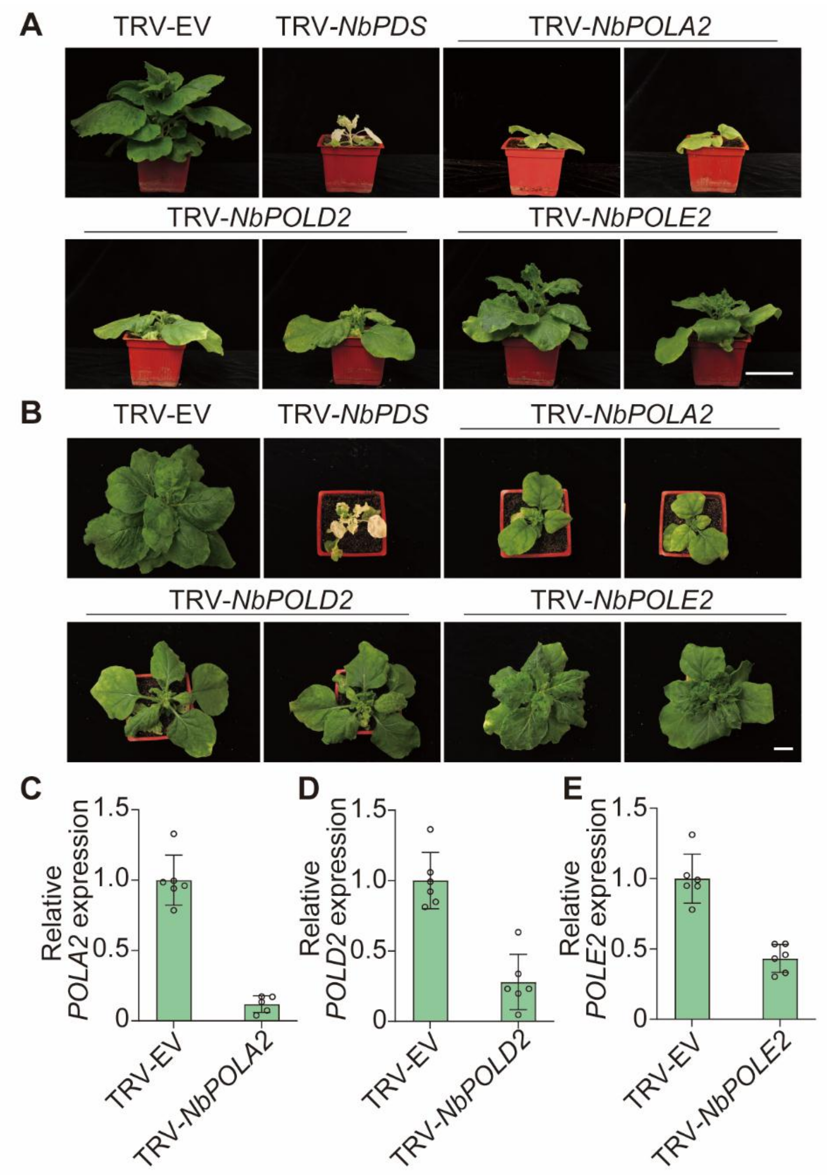


**Figure S3**. **Developmental phenotypes of *NbPOLA2*-, *NbPOLD2*-, and *NbPOLE2*-silenced (TRV-NbPOLA2, TRV-NbPOLD2, and TRV-NbPOLE2, respectively) *N. benthamiana* plants and silencing efficiency.** Lateral (A) and top (B) views are shown. TRV empty vector (TRV-EV) and TRV-PDS are included as controls. Two different constructs were used for silencing of each target gene (see Tables S2 and S3); for each gene, a plant inoculated with version 1 is shown on the left, and a plant inoculated with version 2 is shown on the right. Images were taken at 21 days post-inoculation. Scale bar: 5 cm. C. *NbPOLA2*, *NbPOLD2*, and *NbPOLE2* transcript accumulation in silenced and control plants, measured by qRT-PCR. TRV-NbPOLA2-A, TRV-

NbPOLD2-2, and TRV-NbPOLE2-1 (see Tables S2 and S3) are used in this panel and in all subsequent experiments. *NbActin* was used as reference gene. Values are presented relative to those in the TRV-EV plants. Error bars represent SD with n=6 independent biological replicates. This experiment was repeated twice with similar results.

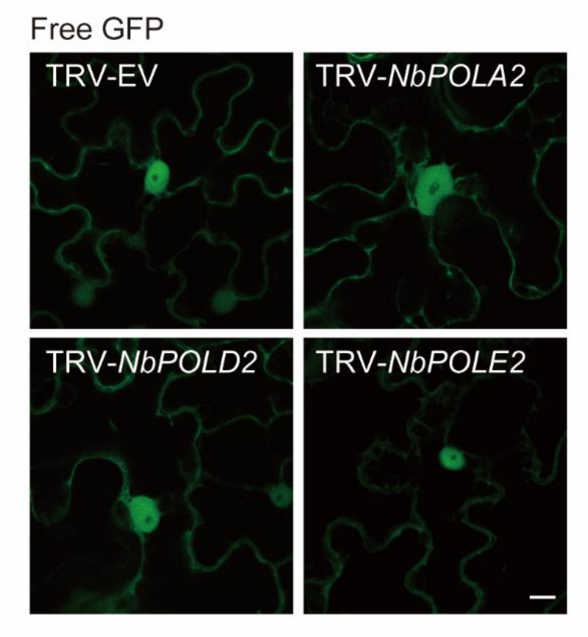


**Figure S4. Silencing of *NbPOLA2*, *NbPOLD2*, or *NbPOLE2* does not affect Agrobacterium-mediated transient expression of free GFP in *N. benthamiana*.** TRV-EV: empty vector control. Scale bar: 10 µm. This experiment was repeated twice with similar results.

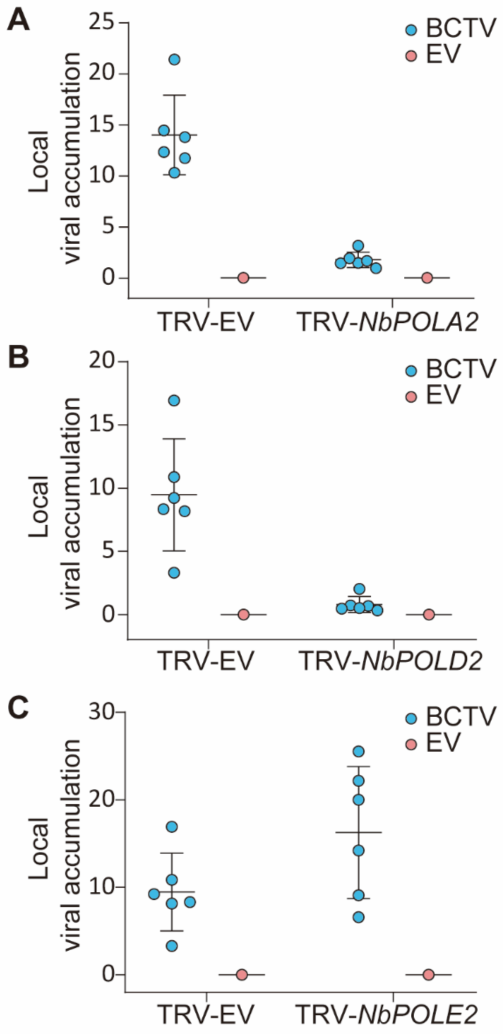


**Figure S5. Viral accumulation in local BCTV infections in *NbPOLA2*-, *NbPOLD2*-, *NbPOLE2*-silenced or control (TRV-EV) *N. benthamiana* plants measured by qPCR.** Plants inoculated with the empty vector (EV) are used as negative control. Error bars represent SD with n=6 independent biological replicates. The 25S ribosomal DNA interspacer (*ITS*) was used as reference gene; values are presented relative to *ITS*. These experiments were repeated three times with similar results.

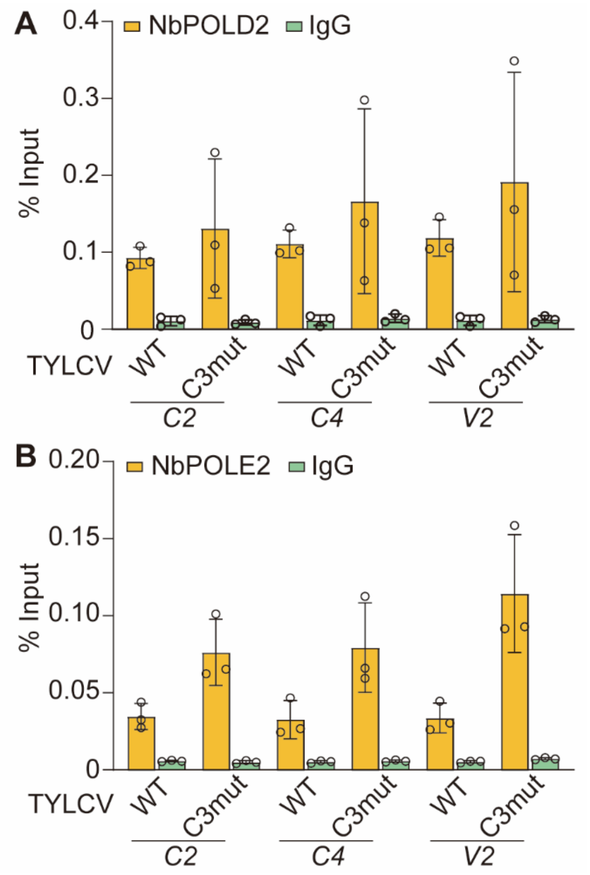


**Figure S6. Binding of NbPOLD2 (A) and NbPOLE2 (B) to the C2, C4, and V2 regions of the TYLCV genome.** Error bars represent SD with n=3 independent biological replicates. These experiments were repeated at least twice with similar results.

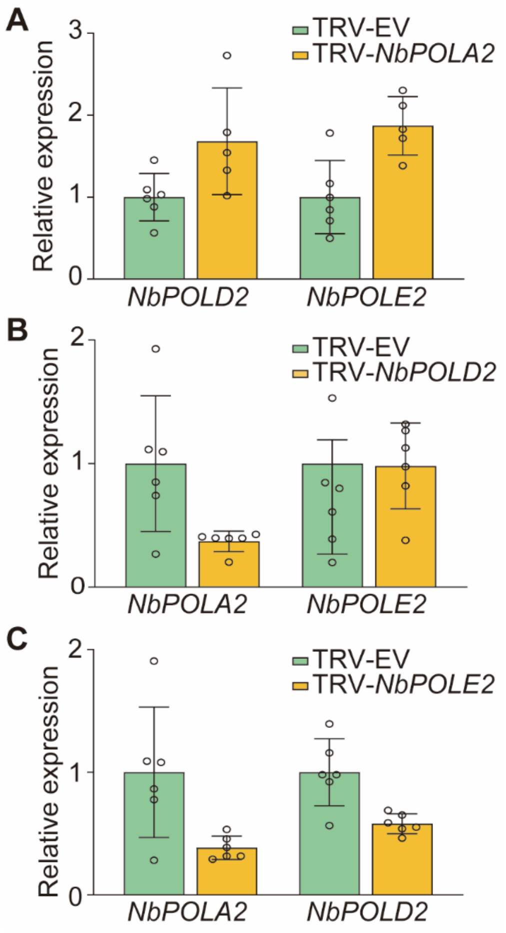


**Figure S7. *NbPOLD2* and *NbPOLE2* transcript accumulation in *NbPOLA2*-silenced plants (A), *NbPOLA2* and *NbPOLE2* transcript accumulation in *NbPOLD2*-silenced plants (B), and *NbPOLA2* and *NbPOLD2* transcript accumulation in *NbPOLE2*-silenced plants (C), measured by qRT-PCR.** *NbActin* was used as reference gene. Error bars represent SD with n=6 independent biological replicates. This experiment was repeated twice with similar results.

| | EV_1 | EV_2 | EV_3 | TYLCV_1 | TYLCV_2 | TYLCV_3 |
|---|---|---|---|---|---|---|
| ***POLA1*** | | | | | | |
| Niben101Scf01802g02001 | 188 | 216 | 310 | 21 | 19 | 13 |
| Niben101Scf04003g04002 | 256 | 257 | 409 | 36 | 22 | 58 |
| ***POLA2*** | | | | | | |
| **Niben101Scf18951g00009** | **182** | **169** | **233** | **200** | **146** | **152** |
| Niben101Scf01073g04022 | 89 | 51 | 83 | 31 | 19 | 19 |
| ***POLA3*** | | | | | | |
| Niben101Scf01950g03015 | 219 | 108 | 206 | 127 | 100 | 92 |
| Niben101Scf00366g01013 | 498 | 415 | 589 | 308 | 336 | 315 |
| ***POLA4*** | | | | | | |
| Niben101Scf13695g01002 | 65 | 66 | 119 | 25 | 24 | 19 |
| Niben101Scf18347g00003 | 45 | 58 | 86 | 17 | 8 | 4 |
| Niben101Scf04053g02011 | 101 | 110 | 148 | 7 | 17 | 13 |
| ***POLD1*** | | | | | | |
| Niben101Scf02230g03027 | 363 | 258 | 421 | 393 | 245 | 323 |
| Niben101Scf00215g00022 | 74 | 63 | 68 | 62 | 30 | 56 |
| Niben101Scf10041g00012 | 85 | 53 | 84 | 77 | 35 | 51 |
| ***POLD2*** | | | | | | |
| Niben101Scf02793g17003 | 50 | 42 | 42 | 45 | 34 | 26 |
| Niben101Scf02793g17022 | 66 | 43 | 65 | 44 | 56 | 38 |
| **Niben101Scf07121g02003** | **311** | **264** | **362** | **442** | **347** | **341** |
| Niben101Scf14412g00012 | 0 | 0 | 0 | 0 | 0 | 0 |
| Niben101Scf01230g04020 | 0 | 0 | 0 | 0 | 0 | 0 |
| Niben101Scf09867g02025 | 0 | 0 | 0 | 0 | 0 | 0 |
| Niben101Scf06725g02018 | 0 | 0 | 0 | 0 | 0 | 0 |
| Niben101Scf02449g02012 | 0 | 0 | 0 | 0 | 0 | 0 |
| Niben101Scf08523g00014 | 0 | 0 | 0 | 0 | 0 | 0 |
| Niben101Scf07590g05005 | 0 | 0 | 0 | 0 | 0 | 1 |
| ***POLD3*** | | | | | | |
| Niben101Scf00160g10002 | 179 | 125 | 185 | 157 | 193 | 142 |
| ***POLD4*** | | | | | | |
| Niben101Scf02887g01014 | 6 | 11 | 13 | 2 | 2 | 3 |
| Niben101Scf00171g00003 | 65 | 51 | 79 | 72 | 71 | 70 |
| Niben101Scf08228g04010 | 133 | 99 | 117 | 128 | 162 | 154 |
| ***POLE1*** | | | | | | |
| Niben101Scf11937g01021 | 1056 | 610 | 1192 | 937 | 383 | 783 |
| ***POLE2*** | | | | | | |
| **Niben101Scf08137g03002** | **550** | **421** | **717** | **187** | **231** | **207** |
| Niben101Scf00887g00009 | 138 | 103 | 152 | 68 | 47 | 46 |
| Niben101Scf02174g02008 | 3 | 0 | 5 | 15 | 1 | 2 |

**Table S1. Orthologues of the Arabidopsis genes encoding the subunits of DNA polymerases α (POLA), δ (POLD), and ε (POLE) in *N. benthamiana*, and their expression in TYLCV-locally infected samples and the corresponding controls.** Data (in reads per million) are from Wu et al., 2019. Three independent biological replicates (1-3) are shown. EV: empty vector. Genes selected for further experiments are indicated in red.

| PLASMIDS | | |
|---|---|---|
| **Expression cassette/virus** | **Source** | **Destination vector** |
| TYLCV-WT | Rosas-Diaz et al., 2018 | pGWB501 |
| TYLCV-C3mut | This paper | pGWB501 |
| pBIN1.2 (BCTV) | Briddon et al., 1989 | - |
| 35S:*C3-GFP* | Wang et al., 2017a | pGWB5 |
| 35S:*GFP-C3* | Wang et al., 2017a | pGWB6 |
| 35S:*SlPOLA2-RFP* | This paper | pGWB554 |
| 35S:*SlPOLA2-GFP* | This paper | pGWB505 |
| 35S:*NbPOLA2-RFP* | This paper | pGWB554 |
| 35S:*NbPOLA2-GFP* | This paper | pGWB505 |
| 35S:*NbPOLD2-RFP* | This paper | pGWB554 |
| 35S:*NbPOLD2-GFP* | This paper | pGWB505 |
| 35S:*NbPOLE2-RFP* | This paper | pGWB554 |
| 35S:*NbPOLE2-GFP* | This paper | pGWB505 |
| TRV2:*NbPOLA2*-1 | This paper | pTRV2 |
| TRV2:*NbPOLA2*-2 | This paper | pTRV2 |
| TRV2:*NbPOLD2*-1 | This paper | pTRV2 |
| TRV2:*NbPOLD2*-2 | This paper | pTRV2 |
| TRV2:*NbPOLE2*-1 | This paper | pTRV2 |
| TRV2:*NbPOLE2*-2 | This paper | pTRV2 |
| 35S:*C3-TYLCV-YN* | This paper | pGTQL1211YN |
| 35S:*C3-BCTV-YN* | This paper | pGTQL1211YN |
| 35S:*C3-TGMV-YN* | This paper | pGTQL1211YN |
| 35S:*SlPOLA2-YC* | This paper | pGTQL1221YC |
| 35S:*NbPOLA2-YC* | This paper | pGTQL1221YC |
| 35S:*NbPOLD2-YC* | This paper | pGTQL1221YC |
| 35S:*NbPOLE2-YC* | This paper | pGTQL1221YC |
| AD-*SlPOLA2* | This paper | pGADT7 |
| BD-*C3* (TYLCV) | This paper | pGBKT7 |

**Table S2. Plasmids and constructs used in this work.**

| AMPLIFICATION TARGET | SOURCE | SEQUENCE 5’ ~ 3’ |
|---|---|---|
| **Oligonucleotides to clone in pENTR™/D-TOPO® (F: CACC) and the pDONR™-zeo entry vector (F: GGGGACAAGTTTGTACAAAAAAGCAGGCTNN, R: GGGGACCACTTTGTACAAGAAAGCTGGGTN) entry vector (Thermo Scientific)** | | |
| TOPO-*C3* (TYLCV) (with stop codon) | This paper | F: CACCATGGATTCACGCACAG<br>R: TTAATAAAATTTATATT |
| TOPO-*C3* (TYLCV) (without stop codon) | Medina-Puche et al., 2019 | F: CACCATGGATTCACGCACAG<br>R: ATAAAATTTATATTTTATATC |
| TOPO-*C3* (BCTV) (without stop codon) | This paper | F: CACCATGAATGTAATAGAGGA<br>R: GTACAAGTTCATTGCAACAC |
| TOPO-*SlPOLA2* (without stop codon) | This paper | F: CACCATGGAAGAGGAAATCAAAGC<br>R: TATACGAAGGACTGAAGCAC |
| pDONR-zeo-*NbPOLA2* (without stop codon) | This paper | F:GGGGACAAGTTTGTACAAAAAAGCAGGCTTCATGGAAGAGCAAATCAAAGCTGA<br>R:GGGGACCACTTTGTACAAGAAAGCTGGGTCTATACGAATAACTGAAGCACTTGACAAATCAC |
| pDONR-zeo-*NbPOLD2* (without stop codon) | This paper | F:GGGGACAAGTTTGTACAAAAAAGCAGGCTTCATGAGTTCAGAATTTGATTTTCTCCT<br>R:GGGGACCACTTTGTACAAGAAAGCTGGGTCTGAGTGGATTTGAGTAGCAAAGCTG |
| pDONR-zeo-*NbPOLE2* (without stop codon) | This paper | F:GGGGACAAGTTTGTACAAAAAAGCAGGCTTCATGCTGTGTATCCATCACCTGCAGTAC<br>R:GGGGACCACTTTGTACAAGAAAGCTGGGTCCAATGCTGAGAGTTCTACTTCCTGA |
| **Oligonucleotides to clone in pTRV2 entry vector** | | |
| TRV2:*NbPOLA2*-1 | This paper | F: ATCGGAATTCAGGAGAACCCCAATGATG<br>R: ATCGGAGCTCCGCTGCTTTGCAGTAAAA |
| TRV2:*NbPOLA2*-2 | This paper | F: ATCGGAATTCATGAGGTGAAAGTGGCTTGC<br>R: ATCGGAGCTCAAAAGCGTGAAGCACTTGA |
| TRV2:*NbPOLD2*-1 | This paper | F: ATCGGAATTCTCCAGATTGCAGCAAGTATAC<br>R: ATCGGAGCTCCACCACTGCCACTCCAGTATCTG |
| TRV2:*NbPOLD2*-2 | This paper | F: ATCGGAATTCCGCCCTGCCACAGCAGCCTCTC<br>R: ATCGGAGCTCCTGAGTCTAATGGATATCTTTTGC |
| TRV2:*NbPOLE2*-1 | This paper | F: ATCGGAATTCATGCTGTGTATCCATCACCTG<br>R: ATCGGAGCTCCTTGCCTGAGACTTGAAATTGA |
| TRV2:*NbPOLE2*-2 | This paper | F: ATCGGAATTCAGAAGACTTTGGGAAATCTG<br>R: ATCGGAGCTCGCAGAGATGGCTCTGATGTAT |
| **Oligonucleotides to clone in Y2H vectors** | | |
| AD-*SlPOLA2* | This paper | F: GCCAGTGAATTCATGGAAGAGGAAATCAAAG<br>R: GCTCGATGGATCCTTATATACGAAGGACTG |
| BD-*C3* (TYLCV) | This paper | F: GAGGCCGAATTCATGGATTCACGCAC<br>R: CAGGTCGACGGATCCTTAATAAAATTTATATTTTATATCATG |
| **Oligonucleotides for gene expression** | | |
| *25S ribosomal DNA interspacer (ITS)* | Rosas-Diaz et al., 2018 | F: ATAACCGCATCAGGTCTCCA<br>R: CCGAAGTTACGGATCCATTT |
| *NbACTIN* | Maimbo et al., 2010 | F: CGGAATCCACGAGACTACATAC<br>R: GGGAAGCCAAGATAGAGC |
| *Rep* (TYLCV) | Rosas-Diaz et al., 2018 | F: TGAGAACGTCGTGTCTTCCG<br>R: TGACGTTGTACCACGCATCA |
| *C2* (TYLCV) | Wang et al., 2017b | F: ACCTTCGTCACCCTCTACGA<br>R: AAACGCCATTCTCTGCCTGA |
| *C3* (TYLCV) | This paper | F: TGGACGACATTACAGCCTCA<br>R: ACAATACATGATCAACTGCTCTGA |
| *C4* (TYLCV) | Rosas-Diaz et al., 2018 | F: TGCTGACCTCCTCTAGCTGA<br>R: ATCCGAACATTCAGGCAGCT |
| *CP* (TYLCV) | Wang et al., 2017b | F: TGGAAGCAGCCCAATGGATT<br>R: GTTCTCGTACTTGGCTGCCT |
| *V2* (TYLCV) | Wang et al., 2017b | F: ATCTGTTGTAAGGGCCCGTG<br>R: CTTTCGGTACATGGGCCTGT |
| *IR* (TYLCV) | This paper | F: GGCATGTTGAAATGAATCGG<br>R: GGTCCCACATATTGCAAGAC |
| *Rep* (BCTV) | This paper | F: AATGCAAGAATGGGCTGATGC<br>R: GCCCACATAGTCTTCCCTGTT |
| *NbPOLA2* | This paper | F: TGGAGAAAGAAGTGAAGGGGAAG |

| | | |
|---|---|---|
| | | R: TGACAAATCACAGCTTCCGTG |
| *NbPOLD2* | This paper | F: TTGATTTTCTCCTTTTCCAGATTGC<br>R:TTTGCTGGATCACTGGACCC |
| *NbPOLE2* | This paper | F: ACTATTCCGCCAAGATTTGCT<br>R: TGGTTTCTTCTGTTGAGGGAGG |
| **Oligonucleotides for testing virion and complementary viral stand** | | |
| *TAG* | Rodriguez-Negrete et al., 2014 | AGTTTAAGAACCCTTCCCGC |
| *OCS* | Rodriguez-Negrete et al., 2014 | GGACTTTACATGGGCCTTCAC |
| *OVS* | Rodriguez-Negrete et al., 2014 | GAAGGCTGAACTTCGACAGC |
| *OCS-TAG* | Rodriguez-Negrete et al., 2014 | AGTTTAAGAACCCTTCCCGCGGACTTTACATGGGCCTTCAC |
| *OVS-TAG* | Rodriguez-Negrete et al., 2014 | AGTTTAAGAACCCTTCCCGCGAAGGCTGAACTTCGACAGC |
| **Oligonucleotides for generating TYLCV-C3mut** | | |
| TYLCV-C3mut | This paper | F: GGGGATTGTTTATCTCCTAAATAAAAACGCCATTCTC<br>R: GAGAATGGCGTTTTTATTTAGGAGATAAACAATCCCC |

### Table S3. Primers used in this work.
